## Supplementary material for "Physically asymmetric division of the *C. elegans* zygote ensures invariably successful embryogenesis": Figure 1 - supplement 1 - source code: Figure 1 - supplement 1 -source code.R

```
##run in R version 3.6.1

library(openxlsx)
library(beeswarm)

# function to plot correlation with confidence intervals
plot.corr <- function (filename, data, xvar, yvar, xlab="x", ylab="y", groupvar="Group",title="correlation", plot.width=2.5, save=T, leg=F, xlim=F, ylim=F){
  data <- as.data.frame(data)
  
  if(save==T){ 
    pdf(filename,width = plot.width, height = plot.width*1.05, pointsize = 10, family='Helvetica', useDingbats = FALSE)
    par(mar=c(2.3,2.3,1,0.2), mgp=c(1.25,0.5,0), cex.lab=0.8, cex.axis=0.7, cex.main=0.8, tcl=-.35)
  }
  
  v1 <- data[,xvar]
  v2 <- data[,yvar]
  g <- factor(data[,groupvar])
  
  xlim <- if(all(xlim==F)) xlim <- range(v1, na.rm = T)
  ylim <- if(all(ylim==F)) ylim <- range(v2, na.rm = T)
  
  plot(v2~v1, col=pal[g], pch=20, xlab=xlab, ylab=ylab, main=title, xlim=xlim, ylim=ylim)
  
  m <- lm(v2~v1)
  ct <- cor.test(v2,v1, method = "pearson")

    abline(m, lwd=1.5)
    
    newx <- seq(min(v1, na.rm = T)*0.9,max(v1, na.rm = T)*1.1, length.out=length(v1))
    prd <- predict.lm(m,newdata=data.frame(v1=newx),interval = c("confidence"),level = 0.90, type="response")
    lines(newx,prd[,2],col="grey",lty=2)
    lines(newx,prd[,3],col="grey",lty=2)

  mtext(paste(c("r = ",round(unlist(ct[c("estimate")]), digits=2),", p = ", signif(ct$p.value, digits=-1)), collapse = ""), side=3, cex=0.7, line=-1.2) #adj=0.1, padj=0, )
  if(leg==T)legend("bottomright",levels(g),pch=16, cex=0.8, col=pal[factor(levels(g))], inset=0.02)
  
  if(save==T) dev.off()
}

# Volumes of embryos measured in 3D and 2D ####
setwd("/Users/jankele/UPGON/LIN-5/MANUSCRIPT/New_submission/Figure 1 - supplement/")
pal<-c("#2E5268", "#64A8A1", "#D19812", "#C93230", "azure4") # colore pallete

# Panel A - technical error of 2D manual measurement ####
# read data with repeated measurements of cell sizes (sd already calcuted beforehand)
dt <- read.xlsx("Figure 1 - supplement 1 - source data 1.xlsx", rows = 1:32)
dt <- dt[dt$n > 2,] #some embryos were measured only twice, keep only those measured more than twice
dt[,c("AB.rel","sd","vc")] <- dt[,c("AB.rel","sd","vc")]*100 #express in percent
boxplot(dt$vc, ylab="Technical error")
beeswarm(dt$vc, add=T)

# Panel C-D ####
Embs <- read.xlsx("Figure 1 - supplement 1 - source data 1.xlsx", sheet = 2)
Embs[,c("AB.rel","AB.3D.rel")] <- Embs[,c("AB.rel","AB.3D.rel")]*100 # in percent of size
Embs$Group <- factor(Embs$Group, levels=c("wt","ctrl","alive","dead","inverted"))
Embs$Tot.Volume <- Embs$AB.3D+Embs$P1.3D

plot.corr("AB_size_2D_vs_3D.pdf", Embs, "AB.rel", "AB.3D.rel",
          title="Relative size of AB cell", xlab="2D estimation", ylab="3D measurement", groupvar = "Group",
          plot.width = 1.75,
          save=F,
          leg=T
          )

# Relative AB size estimation accuracy ####
EstimationError <- (Embs$AB.rel-Embs$AB.3D.rel)
t.test(EstimationError) #p = 0.02, -0.33%
sd(EstimationError, na.rm = T)

  #not shown in the manuscript
  boxplot(EstimationError, main="AB size estimation error", ylab="2D estimation - 3D measurement %"); abline(h=0, lty=2)
  beeswarm(EstimationError, add=T, cex=0.8)

# correlation between embryo size and estimation error between 2D and 3D
df <- Embs
df$error <- EstimationError

# remove outliers for estimation error before plotting correlation
df <- df[-which(EstimationError==boxplot(EstimationError, plot=F)$out),]


#no correlation between error and relative AB volume
plot.corr("AB_size_corr_3D_error.pdf", df, "AB.3D.rel", "error",
          title="Estimation Error ~ AB volume %", 
          xlab="relative AB volume", 
          ylab="2D - 3D error", 
          groupvar = "Group",
          plot.width = 1.75,
          save=T
)
#r=-0.26, p=0.1 

#no correlation between error and total volume (not shown in the manuscript)
plot.corr("file", df, "Tot.Volume", "error",
          title="Estimation Error ~ total volume", 
          xlab="Tot embryo volume", 
          ylab="2D - 3D error", 
          groupvar = "Group",
          plot.width = 1.75,
          save=F
)
#r=0.3, p=0.17
```
