## Supplementary material for "Physically asymmetric division of the *C. elegans* zygote ensures invariably successful embryogenesis": Figure 1 - supplement 2 - source code: Figure 1 - supplement 2 - source code.R

```
library(ggpubr)
library(openxlsx)

#color pallete
pal<-c("#2E5268", "#64A8A1", "#D19812", "#C93230")

# PAR-2 analysis ####
setwd("/Users/jankele/UPGON/LIN-5/MANUSCRIPT/Revisions/Source-Code_and_Data/Figure 1 - supplement 2/") #Change to source data file destination

#load annotation
annot <- read.xlsx("Figure 1 - supplement 2 - source data.xlsx", sheet=1)
rownames(annot) <- annot$ID

#load raw data
tab <- read.xlsx("Figure 1 - supplement 2 - source data.xlsx", sheet=2)
tab <- tab[tab$cell!="bg",]

#remove camera background intensity measured for each embryo
tab[,c(4:6)] <- tab[,c(4:6)] - annot[tab$embryo,"background"]

# aggregate intensity data (min, max) by embryo and cell
tab <- aggregate(tab[,c(4:6)], by=list(tab$embryo, tab$cell), mean)
colnames(tab)[1:2] <- c("ID","Cell")
tab <- tab[order(tab$ID, tab$Cell),]

#merge it with annot
tab <- merge(tab, annot, by="ID")
tab$CellSize <- ifelse(tab$Cell=="AB", tab$AB.size, tab$P1.size)

#approximate total intensity by multiplying area by mean pixel intensity for respective cell
tab$CellInt <- tab$Mean*tab$CellSize
totInt <- aggregate(tab$CellInt, by=list(tab$ID), sum)
rownames(totInt) <- totInt$Group.1
tab$totInt <- totInt[tab$ID,"x"] 

tab$AB.rel <- tab$AB.rel*100
tab$Exp[tab$Exp=="Upshift"] <- "Equalized"
tab$Exp <- factor(tab$Exp)

#calculate intensity ratio between P1 and AB
P1ratio <- aggregate(tab$Mean, by=list(tab$ID), function(x)x[2]/x[1])
rownames(P1ratio) <- P1ratio$Group.1
tab$P1ratio <- P1ratio[tab$ID,"x"] 

#different imaging conditions every day of imaging
  aggregate(tab$totInt, list(tab$date), summary)
  #plot only the ratio, absolute values are meaningless

#assuming that total loads of PAR-2 are the same in these worms, we can normalize by total intensity
  #normalize by total intenstity of whole embryo
  tab$norm <- tab$totInt/mean(tab$totInt)
  tab$MeanNorm <- tab$Mean/tab$norm
  #this normalization has no effect on the ratio of the two cells

#Compare groups
  tab$fact <- as.factor(paste0(tab$Cell,"_", tab$Exp))
  
  m <- compare_means(MeanNorm~fact, data=tab, method = "t.test")
  write.xlsx(m,file="P-values_intensity_PAR-2.xlsx",rowNames = F)
  
  #remove non-signif comparisons from the list
  m <- m[m$p.signif!="ns",]
  compr <- split(unlist(m[1:nrow(m),2:3]), seq(nrow(m[1:nrow(m),])))

scp <- ggscatter(tab, "AB.rel", "MeanNorm",
                 color="fact", palette=pal, show.legend = F, size=1,
                 title="PAR-2 intensity ~ AB size",
                 xlab="Relative AB size", 
                 ylab="GFP::PAR-2 intensity [AU]",
                 theme=theme_pubr(base_size = 6, base_family = "Lato", margin =F, legend ="none")
                 )
#not shown (not shown in the manuscript)
scp <- ggpar(scp, legend ="none", legend.title = "")

bxp <- ggboxplot(tab, x = "fact", y ="MeanNorm",
                 title="GFP::PAR-2 intensity",
                 ylab=F, xlab=FALSE,
                 color = "black",
                 add="jitter",
                 palette = pal,
                 add.params=list(color="fact", size=1),
                 show.legend=F,
                 outlier.shape=NA,
                 theme=theme_pubr(base_size = 6, base_family = "Lato", margin =F, legend ="none")
                 )
#  stat_compare_means(comparisons = compr, hide.ns=T, label = "p.signif", show.legend=FALSE) # Add pairwise comparisons p-value
p <- ggarrange(bxp, scp, ncol=2, nrow=1, common.legend = T, legend = "none", align = "h")
p

ggsave(paste0("PAR-2_normalizedIntensity.pdf"),p,width=14, height =8,units="cm")

#plot PAR-2 ratio (not shown in the manuscript)
scatterhist <- ggscatterhist(tab, "AB.rel", "P1ratio",
                             color="Exp", 
                             palette=pal[c(1,4)], 
                             size=2,
                             #shape = "Outcome",
                             margin.plot = "boxplot",
                             xlab="Relative AB size %",
                             theme=theme_pubr(base_size = 8),
                             #legend="none",
                             ylab="P1/AB PAR-2 ratio"
) + stat_compare_means()
scatterhist
ggsave(paste0("PAR-2_Ratio.pdf"),scatterhist, width=8, height =8,units="cm")

t.test(tab$P1ratio~tab$Exp)
#p-value = 0.258
#group Ctrl mean in group Equalized 
#   1.71          1.75

### plot raw Mean values (not shown, but same conclusion)

scp <- ggscatter(tab, "AB.rel", "Mean",
                 color="fact", palette=pal, show.legend = F, size=1,
                 title="PAR-2 intensity ~ AB size",
                 xlab="Relative AB size", 
                 ylab="GFP::PAR-2 intensity [AU]",
                 theme=theme_pubr(base_size = 6, base_family = "Lato", margin =F, legend ="none")
)

scp <- ggpar(scp, legend ="none", legend.title = "")

bxp <- ggboxplot(tab, x = "fact", y ="Mean",
                 title="GFP::PAR-2 intensity",
                 ylab=F, xlab=FALSE,
                 color = "black",
                 add="jitter",
                 palette = pal,
                 add.params=list(color="fact", size=1),
                 show.legend=F,
                 outlier.shape=NA,
                 theme=theme_pubr(base_size = 6, base_family = "Lato", margin =F, legend ="none")
)

p <- ggarrange(bxp, scp, ncol=2, nrow=1, common.legend = T, legend = "none", align = "h")
p
ggsave(paste0("PAR-2_raw_Intensity.pdf"),p,width=14, height =8,units="cm")

# MEX-5 analysis ####
tab <- read.xlsx("Figure 1 - supplement 2 - source data.xlsx", sheet=3)

tab$AB.rel <- tab$AB.rel*100
names(tab)[c(1:2)] <- c("Exp","Size")

#remove camera background intensity = 210 for binning 2
tab[,c("AB.int","P1.int")] <- tab[,c("AB.int","P1.int")] - 210

#calculate back total intensity by multiplying area of each cell by mean pixel intensity for respective cell
tab$total.int <- tab$AB.int*tab$AB.size+tab$P1.int*tab$P1.size
tab$Exp[tab$Exp=="ctrl"] <- "Ctrl"
tab$Exp[tab$Exp=="meta"] <- "Equalized"
tab$Exp <- factor(tab$Exp )

tab$P1ratio <- tab$P1.int/tab$AB.int 

aggregate(tab$total.int, list(tab$Day), summary)
t.test(tab$total.int~tab$Day)
# intensities from the two days are dramatically different
# p-value = 1.567e-06
# a 755728.6  b 1006512.3

#normalize AB and P1 intensity values with respect to the mean intenstity of AB in ctrls from the respective day
AB.mean <- aggregate(tab$AB.int[tab$Exp=="Ctrl"], list(tab$Day[tab$Exp=="Ctrl"]), mean)
#probably different imaging conditions each day

tab$AB.norm <- 0
tab$AB.norm[tab$Day=="a"] <- tab$AB.int[tab$Day=="a"]/AB.mean$x[1]
tab$P1.norm[tab$Day=="a"] <- tab$P1.int[tab$Day=="a"]/AB.mean$x[1]

tab$AB.norm[tab$Day=="b"] <- tab$AB.int[tab$Day=="b"]/AB.mean$x[2]
tab$P1.norm[tab$Day=="b"] <- tab$P1.int[tab$Day=="b"]/AB.mean$x[2]

#transform table into a long format for plotting with ggplot##
tab1 <- tab[,c(1:5,11)]
tab1$Cell <- "AB" 
names(tab1)[5] <- "raw" #AB.int
names(tab1)[6] <- "intensity" #AB.norm

tab2 <- tab[,c(1:4,6,12)]
tab2$Cell <- "P1"
names(tab2)[5] <- "raw"
names(tab2)[6] <- "intensity"

tabL <- rbind(tab1, tab2)
tabL$fact <- as.factor(paste0(tabL$Cell,"_", tabL$Exp))

# difference in MEX-5 ratio between upshifted and control embryos  
t.test(tab$P1ratio~tab$Exp)
#p-value = 8.946e-05
#group Ctrl mean in group Equalized 
#   0.5624167             0.6400186 

#difference: 0.078 = 8%

#SD for each group and for the difference
s <- aggregate(tab$P1ratio, list(tab$Exp), sd); s
#Ctrl 0.0332
#Equalized 0.0344

#SD of difference is SQRT(SD1^2+SD2^2)
s <- s[,2]
sqrt(s[1]^2+s[2]^2) #0.04779

#Compare groups
m <- compare_means(intensity~fact, data=tabL, method = "t.test")
write.xlsx(m,file="P-values_normalized_intensity-MEX-5.xlsx",rowNames = F)
m <- m[m$p.signif!="ns",]

scp <- ggscatter(tabL, "AB.rel", "intensity",
                 color="fact", palette=pal, show.legend = F, size=1,
                 title="MEX-5 intensity ~ AB size",
                 xlab="Relative AB size", ylab="mCherry::MEX-5 intensity [AU]",
                 theme=theme_pubr(base_size = 6, base_family = "Lato", margin =F, legend ="none")
)

scp <- ggpar(scp, legend ="none", legend.title = "")

bxp <- ggboxplot(tabL, x = "fact", y ="intensity",
                 title="mCherry::MEX-5 intensity",
                 ylab=F, xlab=FALSE,
                 color = "black",
                 add="jitter",
                 palette = pal,
                 add.params=list(color="fact", size=1),
                 show.legend=F,
                 outlier.shape=NA,
                 theme=theme_pubr(base_size = 6, base_family = "Lato", margin =F, legend ="none")
)
#  stat_compare_means(comparisons = compr, hide.ns=T, label = "p.signif", show.legend=FALSE) # Add pairwise comparisons p-value

p <- ggarrange(bxp, scp, ncol=2, nrow=1, common.legend = T, legend = "none", align = "h")
p
ggsave(paste0("MEX-5_normalizedIntensity.pdf"),p,width=14, height =8,units="cm")

###### ratio of MEX-5  ####
# not shown in the manuscript
scatterhist <- ggscatterhist(tab, "AB.rel", "P1ratio",
                             color="Exp", 
                             palette=pal[c(1,4)], 
                             size=2,
                             margin.plot = "boxplot",
                             xlab="Relative AB size %",
                             theme=theme_pubr(base_size = 8),
                             legend="none",
                             ylab="P1/AB MEX-5 ratio"
) + stat_compare_means()
scatterhist
ggsave(paste0("MEX-5_Ratio.pdf"),scatterhist, width=8, height =8,units="cm")
```
