## Supplementary material for "Physically asymmetric division of the *C. elegans* zygote ensures invariably successful embryogenesis": Figure 2 - source code: Figure_2_source_code-merged.R

```
##run in R version 3.6.1

library(ggpubr)
library(readxl)
library(beeswarm)

# Functions ####
plot.corr <- function (filename, data, xvar, yvar, xlab, ylab, groupvar,title="correlation", plot.width=2.5, save=T, leg=F, xlim=F){
  data <- as.data.frame(data)
  
  if(save==T){ 
    pdf(paste0(today,"_",filename),width = plot.width, height = plot.width*1.05, pointsize = 10, family='Helvetica', useDingbats = FALSE)
    par(mar=c(2.3,2.3,1,0.2), mgp=c(1.25,0.5,0), cex.lab=0.8, cex.axis=0.7, cex.main=0.8, tcl=-.35)
  }
  
  v1 <- data[,xvar]
  v2 <- data[,yvar]
  g <- data[,groupvar]
  
  xlim <- if(all(xlim==F)) xlim <- range(v1, na.rm = T)
  
  plot(v2~v1, col=pal[g], pch=20, xlab=xlab, ylab=ylab, main=title, xlim=xlim)
  m <- lm(v2~v1)
  ct <- cor.test(v2,v1, method = "pearson")
  
  if(ct$p.value < 0.05){
    abline(m, lwd=1.5)
    
    newx <- seq(min(v1, na.rm = T)*0.9,max(v1, na.rm = T)*1.1, length.out=length(v1))
    prd <- predict.lm(m,newdata=data.frame(v1=newx),interval = c("confidence"),level = 0.90, type="response")
    lines(newx,prd[,2],col="grey",lty=2)
    lines(newx,prd[,3],col="grey",lty=2)
  }
  mtext(paste(c("r =",round(unlist(ct[c("estimate")]), digits=2),", p =", signif(ct$p.value, digits=-1)), collapse = ""), side=3, cex=0.8, line=-1.2) #adj=0.1, padj=0, )
  if(leg==T)legend("bottomright",levels(g),pch=16, cex=0.8, col=pal[factor(levels(g))], inset=0.02)
  
  if(save==T) dev.off()
}
rbind.all.columns <- function(x, y) {
  
  x.diff <- setdiff(colnames(x), colnames(y))
  y.diff <- setdiff(colnames(y), colnames(x))
  
  x[, c(as.character(y.diff))] <- NA
  
  y[, c(as.character(x.diff))] <- NA
  
  return(rbind(x, y))
}

#set ouput directory
outDir <- "/Users/jankele/UPGON/LIN-5/MANUSCRIPT/Revisions/Source-Code_and_Data/Figure 2/"
today <- "201221"

#color pallete for plotting groups
pal <- c("#2E5268", "#1B75B9", "#64A8A1", "#D19812", "#C93230", "azure4")

#### Read source data ####
setwd(outDir)
Surv <- as.data.frame(read_excel("Figure_2_source_data-merged.xlsx", na = c("NA","ND"), sheet=1), stringsAsFactors=T)
rownames(Surv) <- Surv$ID

Surv$Outcome <- factor(Surv$Outcome)
Surv$Exp <- factor(Surv$Exp,levels=c("WT","noUpshift","2C","meta")) # Relevel experiment for better plotting

Surv$group <- as.character(Surv$Exp) # Create a hybrid variable to separate Upshifted Alive and Dead for plotting
Surv$group[Surv$Exp=='meta' & Surv$Outcome=="dead"] <- "Up.Dead"
Surv$group[Surv$Exp=='meta' & Surv$Outcome=="alive"] <- "Up.Alive"
Surv$group <- factor(Surv$group, levels=c("WT","noUpshift","2C","Up.Alive","Up.Dead"))
Surv$lin5 <- factor(ifelse(Surv$Exp=="WT","wt","ev571")) # Create a hybrid variable to separate Upshifted Alive and Dead for plotting


# Mean size of upshifted embryos
with(Surv, aggregate(AB.rel, by=list(Exp), function(x)c(mean(x),sd(x),length(x))))
with(Surv, aggregate(AB.rel, by=list(group), function(x)c(mean(x),sd(x),length(x))))

#### Figure 2A ######
#Numbers of embryos in each group      
table(Surv[,c(4:3)])

g <- ggboxplot(Surv, x="group", y="AB.rel", 
               ylab="AB size %", 
               xlab="Experiment",
               palette = pal[c(1:5)],
               add="jitter",
               color="group",
               add.params=list(size=1, jitter=0.4, shape="Outcome"), 
               outlier.shape=NA,
               ylim=c(40, 64)
               )+ scale_shape_manual(values = c(16,4))+ theme_pubr(base_size = 8)
g <- ggpar(g, legend ="none", legend.title = "")
g

outDir <-paste0(outDir,today)
ggsave(paste0(outDir,"_lin-5(ev571)_AB_size_all_embs.pdf"),plot=g, device="pdf", width=3, height=3)

compare_means(AB.rel~group,Surv, method = "t.test", p.adjust.method="BH")
  aggregate(Surv$AB.rel, by=list(Surv$group), mean)
  aggregate(Surv$AB.rel, by=list(Surv$group), length)
  aggregate(Surv$AB.rel, by=list(Surv$group), sd)

#### Figure 2B ######
  #How lethality depens on the AB size
  br <- c(0.3,seq(0.48, 0.56, 0.02))*100 
  
  #equal survival
  Eq <- Surv[Surv$Exp=="meta",]
  Eq$bin <- cut(Eq$AB.rel, breaks=br, labels=paste(br[2:length(br)],"<", sep=""), include.lowest=T)
  
  tab <- table(Eq$bin, Eq$Outcome)
  freq <- as.data.frame(cbind(prop.table(tab,1)))
  freq$class <- rownames(freq)
  freq$lab <- rowSums(tab)

  xt <- table(Surv[,c(3:4)])
  s <- colSums(xt)
  ctrls <- c(xt[2,1:3])/s[1:3]
  ctrls <- data.frame(dead=ctrls, alive=1-ctrls, class=c("Wt", "noUp", "2C"), lab=s[1:3])
  
  freq <- rbind(freq, ctrls)
  freq$died <- freq$dead*100
  freq$class <- factor(freq$class, levels=c(freq$class[1:(length(freq$class)-3)], "2C","noUp","Wt"))
  
#dotchart
  h <- ggdotchart(freq, x="class", y="died", 
                  xlab="AB size %", 
                  ylab="Embryonic lethality %",
                  label="lab", 
                  show.legend=F, 
                  dot.size=3,
                  font.label = list(size = 8, vjust = 0.5, hjust=-0.6),  
                  rotate=T, 
                  ggtheme = theme_pubr(base_size = 10, base_family = "Lato"),
                  add="segments")
  h 
ggsave(paste0(outDir,"lin-5(ev571)_lethality_dotplot.svg"),plot=h, device="svg", width=2.5, height = 2)

#Uncompressed embryos - 45 um beads ####

lowC <- as.data.frame(read_excel("Figure_2_source_data-1.xlsx", na = c("NA","ND"), sheet=2), stringsAsFactors=T)
lowC$AB.rel <- lowC$AB.rel *100
lowC$Experiment <- as.factor(lowC$Experiment)
lowC$Upshift <- as.factor(lowC$Upshift)
lowC$Outcome <- as.factor(lowC$Outcome)


# lethality in equalized upshifted - somewhat compressed
  table(Eq[Eq$AB.rel>48 & Eq$AB.rel<=53,c(3:4)])
  # 90/(90+84) = 52%

# with low compression
  table(lowC[lowC$AB.rel>48 & lowC$AB.rel<=53,c(2,5)])
  # low compression (45um beads) - 8/(17+8) - 32%
  # low compression (C.slip removed) - 7/(7+13) - 35%

  table(lowC[lowC$AB.rel>48 & lowC$AB.rel<=53,c(5)]) # all combined
  # low compression (C.slip removed) - 15/(30+15) - 33%

  outcomes <- matrix(c(84, 90, 17, 8), nrow=2, ncol=2, byrow = T, dimnames=list(c("compressed","uncompressed"),c("Alive","Dead")))
    fisher.test(outcomes) #p-value = 0.086 >> not significant (same for C.slip removed alone)
  outcomes <- matrix(c(84, 90, 13, 7), nrow=2, ncol=2, byrow = T, dimnames=list(c("compressed","uncompressed"),c("Alive","Dead")))
    fisher.test(outcomes) #p-value = 0.086 >> not significant (same for C.slip removed alone)
  
  outcomes <- matrix(c(84, 90, 30, 15), nrow=2, ncol=2, byrow = T, dimnames=list(c("compressed","uncompressed"),c("Alive","Dead")))
  fisher.test(outcomes) #p-value = 0.0304 >> compression leads to elevated lethality
    
#considering all embryos between >48% and <56% AB
  table(Eq[Eq$AB.rel>48 & Eq$AB.rel<=56,c(3:4)])
  # lethality 104/(104+128) - 44.8%
  
  table(lowC[(lowC$AB.rel>48 & lowC$AB.rel<=56)&lowC$Upshift=="meta",c(2,5)])
  #Experiment          alive died  lethality
  #45 um beads          29    9     23.7
  #Coverslip removed    13    7     35
 
  table(lowC[(lowC$AB.rel>48 & lowC$AB.rel<=56)&lowC$Upshift=="meta",c(5)])
  # low compression lethality 16/(16+42) - 27.6%
   
  outcomes <- matrix(c(128, 104, 29, 9), nrow=2, ncol=2, byrow = T, dimnames=list(c("compressed","uncompressed"),c("Alive","Dead")))
   fisher.test(outcomes) #p-value = 0.02016 >>  
   #compression leads to elevated lethality in upshifted overall compared to 45 um beads

  outcomes <- matrix(c(128, 104, 42, 16), nrow=2, ncol=2, byrow = T, dimnames=list(c("compressed","uncompressed"),c("Alive","Dead")))
  fisher.test(outcomes) #p-value = 0.01759 >>  compression leads to elevated lethality in upshifted overall
  
  # the AB sizes in both subsets have the same distribution
  x<- Eq$AB.rel[Eq$Exp=="meta"]
  x2 <- lowC$AB.rel[lowC$Upshift=="meta"]
  ks.test(x,x2) 
  #p-value = 0.8812 the two distributions are the same

      plot(ecdf(x), xlim = range(c(x, x2)), pch=1)
      plot(ecdf(x2), add = TRUE, lty = "dashed")

  #low compression survival
  lcEq <- lowC[lowC$Upshift=="meta",c("ID","Outcome","AB.rel")]
  lcEq$bin <- cut(lcEq$AB.rel, breaks=br, labels=paste(br[2:length(br)],"<", sep=""), include.lowest=T)
  
  tab <- table(lcEq$bin, lcEq$Outcome)
  lcfreq <- as.data.frame(cbind(prop.table(tab,1)))
  lcfreq$class <- rownames(lcfreq)
  lcfreq$lab <- rowSums(tab)
  lcfreq$exp <- "lowC"
  
  freq2 <- freq[1:5,1:4]
  freq2$exp <- "compr"
  colnames(freq2)[2] <- "died"
  
  freq2 <- rbind(lcfreq,freq2)
  freq2[,1:2] <- freq2[,1:2]*100
  
  # not shown in the manuscript
  barplot(died~exp+class, data = freq2, beside=T, 
          xlab="AB size",
          ylab="Lethality %",
          col=pal[c(2,6)],
          legend.text=c("Compressed","Uncompressed"))

#eggshell lenght ####
  t.test(length~lin5, data=Surv)
  boxplot(length~lin5, data=Surv)
  
  aggregate(Surv$length, by=list(Surv$lin5), mean, na.rm=T)
  aggregate(Surv$length, by=list(Surv$lin5), function(x) sum(!is.na(x)))
  aggregate(Surv$length, by=list(Surv$lin5), function(x) sd(na.omit(x)))
  
  testsubset <- Surv[Surv$lin5=="ev571",]
  t.test(length~Config.4C, data=testsubset) # with T-config vs all other lin-5(ts) embryos that were scored
  aggregate(testsubset$length, by=list(testsubset$Config.4C), function(x) sum(!is.na(x)))
  aggregate(testsubset$length, by=list(testsubset$Config.4C), function(x) sd(na.omit(x)))

#OPTOGENETICS strain SV2121####
  
  Opto <- as.data.frame(read_excel("Figure_2_source_data-merged.xlsx", na = c("NA","ND"), sheet=3, range="A1:AA72"), stringsAsFactors=T)
  Opto <- Opto[!Opto$exclude==1,] # exclude some embryos - see source data for the reason for each excluded embryo
  Opto <- Opto[,-c(2:4,8:13)] # drop unused columns
  
  names(Opto)[c(2,4,9,16)] <- c("Act","AB.rel","Size","Delay")
  Opto$Outcome <- factor(Opto$Outcome)
  Opto$Act <- factor(Opto$Act, levels=c("None","Anterior"))
  Opto$AB.rel <- Opto$AB.rel*100 # as percents
  
  #Stats in no light controls and in illuminated embryos
  with(Opto, aggregate(AB.rel, by=list(Act), function(x)c(mean(x),sd(x),length(x))))
  
  # what is lethality in 48-53% bin?
  table(Opto[Opto$AB.rel>48 & Opto$AB.rel<=53,c(2:3)])
  table(Eq[Eq$AB.rel>48 & Eq$AB.rel<=53,c(3:4)])
    #13 died, 16 hatched ->> 44.8% compared to:
    #90 died 84 hatched ->> 51.7%% in lin-5(ts)
    
    outcomes <- matrix(c(84, 90, 16, 13), nrow=2, ncol=2, byrow = T, dimnames=list(c("compressed","uncompressed"),c("Alive","Dead")))
    fisher.test(outcomes) #p-value = 0.55 >> not difference between the two conditions
  
  # what is lethality in 48-50% bin?
  table(Opto[Opto$AB.rel>48 & Opto$AB.rel<=50,c(2:3)])
  table(Eq[Eq$AB.rel>48 & Eq$AB.rel<=50,c(3:4)])
    #5 died, 3 hatched ->> 62.5% this is similar to lin-5
    #27 died 9 hatched ->> 75% in lin-5(ts)
  
    outcomes <- matrix(c(9, 27, 3, 5), nrow=2, ncol=2, byrow = T, dimnames=list(c("compressed","uncompressed"),c("Alive","Dead")))
    fisher.test(outcomes) #p-value = 0.66 >> not difference between the two conditions
  
  ### Figure 2C ####
  g <- ggboxplot(Opto, x="Act", y="AB.rel", 
                 ylab="Relative AB size %", 
                 xlab="Experiment",
                 show.legend=F, 
                 palette = c("black","black"),
                 add="jitter",
                 color="#424243",
                 add.params=list(size=1.5, jitter=0.30, shape="Outcome", color="Act"), 
                 outlier.shape=NA,
                 ylim=c(40, 64)
  ) + scale_shape_manual(values = c(16,4)) + theme_pubr(base_size = 10)
  g <- ggpar(g, legend ="none", legend.title = ""); g
  
  ggsave(paste0(outDir,"_Figure_2C_opto_AB_size_in_groups.pdf"),plot=g, device="pdf", width=2.5, height=3)
  
  t.test(AB.rel~Act, data=Opto) #p-value = 2.949e-16, 
    #mean: Anterior = 50.9, 
    #no light = 57.4
    aggregate(Opto$AB.rel, by=list(Opto$Act), length)
    aggregate(Opto$AB.rel, by=list(Opto$Act), sd)
  
  ### Figure 2D ####
  br <- c(30,48,50,52,54,56)
  Opto$bin <- cut(Opto$AB.rel, breaks=br, labels=paste(br[2:length(br)],"<", sep=""), include.lowest=T)
  
  activated <- Opto[Opto$Act=="Anterior",]
  
  tab <- table(activated$bin, activated$Outcome)
  freq <- as.data.frame(cbind(prop.table(tab,1)))
  freq$class <- rownames(freq)
  freq$lab <- tab[1:nrow(freq),1]+tab[1:nrow(freq),2]
  
  #controls
  table(Opto[Opto$Act=="None",c("Outcome")])
  
  controls <- data.frame(Alive=10/13, Dead=3/13, class="Control", lab=13)
  freq <- rbind(controls, freq)
  freq[,1:2] <- freq[,1:2]*100
  
  h <- ggdotchart(freq, x="class", y="Dead", 
                  xlab="AB size %", 
                  ylab="Embryonic lethality %",
                  label="lab",
                  show.legend=F, 
                  dot.size=3,
                  font.label = list(size = 8, vjust = 0.5, hjust=-1),  
                  sorting = NULL,
                  rotate=T, 
                  ggtheme = theme_pubr(base_size = 10),
                  add="segments")
  h 
  
  ggsave(paste0(outDir,"_Figure_2D_opto_lethality.pdf"),plot=h, device="pdf", width=4, height=3)
  
  
  #P1-AB asynchrony####
  ### Figure 2E ####
  pal <- c("#D19812", "#C93230")
  timing <- as.data.frame(read_excel("Figure_2_source_data-merged.xlsx", na = c("NA","ND"), sheet=4), stringsAsFactors=T)
  timing$Lin_5 <- "ev571"
  
  timing$Emb.size <- timing$AB.size+timing$P1.size
  timing$AB.rel <- (timing$AB.size/timing$Emb.size)*100
  timing$Outcome <- factor(timing$Outcome) #relevel
  
  # plot delay between AB division and P1
  plot.corr("Figure_2E_P1_mitotic_delay.pdf",timing,"AB.rel","Delay","AB size %", "delta P1-AB time [s]", "Outcome", 
            title="Mitotic Delay" , save=F, xlim=c(40,60), leg=T)
  #r=0.81, p=3e-09
  
  # plot cell cycle ratio (not shown)
    # only embryos for which the entire second cell cycle was measured are shown
    ids <- which(!is.na(timing$CC.ratio))
    plot.corr("cell_cycle_ratio.pdf",timing[ids,],"AB.rel","CC.ratio","AB size %", "P1/AB cell cycle ratio", "Outcome", 
              title="Cel cycle ratio" ,save=F, xlim=c(40,60) ,leg=T)
    
    
    # load data from "UV" slowdown experiment ####
    Sl <- as.data.frame(read_excel("Figure_2_source_data-merged.xlsx", na=c("NA","ND"), sheet=5), stringsAsFactors=T)
    Sl <- Sl[!Sl$exclude==1,] #some embryos were filtered either due to initially sick morphology at the one cell stage or due to 
    
    Sl$UV <- "405nm"
    Sl$UV[Sl$UV_sec==0] <- "none"
    
    Sl$AB.rel <- as.numeric(Sl$AB.rel)*100
    Sl$Outcome <- factor(Sl$Outcome)
    
    # create groups to match those in the Sl table ####
    timing$UV <- "none"
    timing$UV_sec <- 0
    
    timing$Experiment <- "timing"
    timing$Upshift <- "ctrl"
    timing$Upshift[grep("M",timing$ID)] <- "meta" # embryos with ID ending M were upshifted in metaphase during first division
    
    # merge tables
    timing <- rbind.all.columns(timing, Sl[,c(intersect(colnames(timing), colnames(Sl)),"AB2.P2","AB2.EMS")] )
    
    timing$Upshift[timing$Upshift=="no"] <- "ctrl" #relabel non-upshifted (mostly wildtype) as controls (plotted separately)
    timing$Upshift <- factor(timing$Upshift)
    timing$Lin_5 <- factor(timing$Lin_5)
    
    # plot delay data in bins for controls and for UV treated embryos and visualise lethality ####
    timing$fact <- "d"
    timing$fact[timing$AB.rel<=48] <- "b"
    timing$fact[timing$AB.rel>48] <- "a"
    timing$fact[timing$AB.rel>=53] <- "c"
    
    timing$fact <- as.factor(timing$fact)
    
    #remove partially equalized embryos with AB>53%, and analyze only upshifted once
    timing <- timing[!(timing$fact=="c"&timing$Upshift=="meta"),]
    
    #plot boxplots
    bp <- timing[,c("ID","AB.rel","Outcome","Delay","Lin_5","fact","UV","UV_sec","Upshift","Emb.size","AB2.P2","AB2.EMS")]
    bp <- bp[!is.na(bp$Delay),]
    bp$UV <- factor(bp$UV, levels = c("none","405nm"))
    bp$Lin_5 <- factor(bp$Lin_5, levels = c("wildtype","ev571"))
    
    # Figure 2F ####
    pdf(paste0(today,"_P1_delay_405nm_slowdown.pdf"),width = 2.5, height = 2.2, pointsize = 10, family='Helvetica', useDingbats = T)
    {
      par(mar=c(2.3,2.3,1,0.2), mgp=c(1.34,0.5,0), cex.lab=0.8, cex.axis=0.7, cex.main=0.8, tcl=-.35,las=3)
      
      s <- bp$Upshift=="ctrl"
      boxplot(Delay~UV+Lin_5,data=bp[s,],
              xlim=c(0.8,8.5),
              ylim=c(0,520),
              xlab="",
              ylab="AB-P1 mitotic delay [sec]",
              col=c("#ffffff","#429DDB"),
              cex.axis=0.6,
              cex.lab=0.6,
              lty=1,
              lwd=0.5,
              outline=F,
              staplewex=0
      )
      
      beeswarm(Delay~UV+Lin_5,data=bp[s,],
               pwpch=c(16,4)[bp$Outcome[s]],
               cex=0.6,
               spacing=1.05,
               corralWidth=1.5,
               corral="random",
               add=T)
      
      bpeq <- bp[bp$Upshift=="meta" & bp$fact%in%c("b","a"),]
      bpeq$fact <- factor(bpeq$fact)
      
      boxplot(Delay~UV+fact,data=bpeq, 
              col=rep(c("#ffffff","#429DDB"),3),
              axes=F,
              add=T,
              at=5:8,
              las=2,
              lty=1,
              lwd=0.5,
              outline=F,
              staplewex=0
      )
      
      beeswarm(Delay~UV+fact,
               data=bpeq,
               pwpch=c(16,4)[bpeq$Outcome],
               lwd=2, 
               spacing=1.2,
               corral="random",
               corralWidth=1.2,
               cex=0.6, 
               add=T, 
               axes=F,
               at=5:8)
      
      #Add %lethality
      
      a<-aggregate(ID~Lin_5+UV+Outcome+fact,data=bp[bp$Outcome=="alive",],length)
      b<-aggregate(ID~Lin_5+UV+Outcome+fact,data=bp[bp$Outcome=="dead",],length)
      
      lethal <- merge(a,b, by=c("Lin_5","UV","fact"), all = T)
      lethal <- lethal[,c(1,2,3,5,7)]
      colnames(lethal)[4:5] <- c("alive","dead")
      lethal[is.na(lethal)] <- 0
      lethal$total <- lethal$alive+lethal$dead
      lethal$lethality <- round(with(lethal, dead/total)*100, digits =0)
      lethal <- lethal[c(1,2,5,8,3,6,4,7),]
      
      #label lethality
      lethal$max <-  aggregate(Delay~Lin_5+fact+UV,data=bp,max)$Delay[c(3,7,4,8,1,5,2,6)]
      text(1:8,lethal$max,labels=paste0(lethal$lethality,"%"), cex=0.6, pos=3, offset = c(0,30))
      
      dev.off()
    }
    
    aggregate(AB.rel~Lin_5+UV+Upshift,data=bp,mean)
    aggregate(AB.rel~fact+Lin_5+UV,data=bp,mean)
    aggregate(AB.rel~fact+Lin_5+UV,data=bp,length)
    aggregate(AB.rel~fact+Lin_5+UV,data=bp,sd)
    
    m <- aggregate(Delay~fact+Lin_5+UV,data=bp,mean)
    aggregate(Delay~fact+Lin_5+UV,data=bp,sd)
    
    # Delay in the later timepoints - not shown####
       #to asses how the slowdown of P1 propagates to its descendants
        
        aggregate(AB2.EMS~Lin_5+Outcome+UV+fact,data=timing[!is.na(timing$Delay),],length)
        aggregate(AB2.EMS~Lin_5+UV+fact,data=timing[!is.na(timing$Delay),],length)
        
        boxplot(AB2.EMS~UV+fact,data=bp,xlim=c(0.8,10.25),ylim=c(200,1200), 
                main="Propagation of the delay after 405 nm P1 slow-down  \n EMS and P2 division delay relative to AB2 division",
                ylab="Division delay relative to AB2 division [sec]",
                col=rep(c("#FFCC66","#ffffff"),4),
                border="#666666",
                at=c(1,0,2,0,3,0,4,5)
        )
        beeswarm(AB2.EMS~UV+fact,data=bp,pwpch=c(13,16)[bp$Outcome],
                 #pwcol=c("firebrick", "black")[bp$Outcome],
                 lwd=2, spacing=1.1,cex=1, add=T, axes=F,at=c(1,0,2,0,3,0,4,5))
        
        
        boxplot(AB2.P2~UV+fact,data=bp,
                col=rep(c("#99CC99","#ffffff"),4),
                border="#666666",
                at=c(6,0,7,0,8,0,9,10),
                add=T
        )
        beeswarm(AB2.P2~UV+fact,data=bp,pwpch=c(13,16)[bp$Outcome],
                 lwd=2, spacing=1.1,cex=1, add=T, axes=F,
                 at=c(6,0,7,0,8,0,9,10)
        )
    
```
