## Supplementary material for "Physically asymmetric division of the *C. elegans* zygote ensures invariably successful embryogenesis": Figure 5 - source code: Figure 5 - source code.R

```
#analysis performed in R version 3.6.1

library(beeswarm)
library(ggpubr)
library(openxlsx)

today <- format(Sys.Date(), "%y%m%d")
pal <- c("#64A8A1", "#D19812", "#C93230", "#838B8B")
#### Functions ####

plot.bxp <-
  function (filename,
            data,
            var1,
            ylab1,
            plot.width = 1.5,
            saveit = T,
            marginal = F,
            ...) {
    if (saveit & !marginal)
      pdf(
        paste0(today, filename),
        width = plot.width,
        height = 2.1,
        pointsize = 10,
        useDingbats = F
      )
    if(!marginal) par(
      mar = c(2.5, 2.5, 0.1, 0.15),
      mgp = c(1.25, 0.5, 0),
      cex.lab = 0.7,
      cex.axis = 0.6,
      cex.main = 0.8,
      tcl = -.35,
      las = 3
    )
    
    v1 <- data[, var1]
    g <- data$Group
    if(!marginal) {
      boxplot(
      v1 ~ g,
      lwd = 0.5,
      outlwd = 1,
      staplewex = 0,
      whisklty = 1,
      outpch = 16,
      ylab = ylab1,
      xlab = NA,
      ...
      )
      beeswarm(
        v1 ~ g,
        add = T,
        pwcol = pal[g],
        pch = 16,
        cex = .6,
        corral = "random",
        corralWidth = 1.15 ,
        spacing = 1
      )
      
    } else {
      boxplot(
        v1 ~ g,
        lwd = 0.5,
        outlwd = 1,
        staplewex = 0,
        whisklty = 1,
        outpch = 1,
        cex=0.4,
        ylab = NA,
        xlab = NA,
        border = pal,
        medlwd = 2,
        axes=F,
        ...
      )
    }
    if (saveit) dev.off()
  }
compare.bxp <-
  function (filename,
            data,
            var1,
            var2,
            ylab1,
            ylab2,
            ylim = F,
            plot.width = 2) {
    pdf(
      paste0(today, filename),
      width = plot.width,
      height = 2.5,
      pointsize = 10,
      useDingbats = F
    )
    par(
      mar = c(2, 2.3, 0.2, 0.15),
      mgp = c(1.25, 0.5, 0),
      cex.lab = 0.7,
      cex.axis = 0.6,
      cex.main = 0.8,
      tcl = -.35,
      las = 3
    )
    par(mfrow = c(1, 2))
    
    v1 <- data[, var1]
    v2 <- data[, var2]
    
    ylim <-
      if (all(ylim == F))
        ylim <- range(c(v1, v2), na.rm = T)
    else
      ylim = ylim
    
    g <- data$Group
    boxplot(
      v1 ~ g,
      lwd = 0.5,
      outlwd = 1,
      staplewex = 0,
      whisklty = 1,
      outpch = NA,
      ylab = ylab1,
      ylim = ylim,
      xlab = NA
    )
    beeswarm(
      v1 ~ g,
      add = T,
      col = pal,
      pch = 16,
      cex = .8,
      corral = "random",
      corralWidth = 1.2
    )
    
    boxplot(
      v2 ~ g,
      lwd = 0.5,
      outlwd = 1,
      staplewex = 0.25,
      whisklty = 1,
      outpch = NA,
      ylab = ylab2,
      ylim = ylim,
      xlab = NA
    )
    beeswarm(
      v2 ~ g,
      add = T,
      col = pal,
      pch = 16,
      cex = .8,
      corral = "random",
      corralWidth = 1.2
    )
    dev.off()
  }
plot.corr <-
  function (filename,
            data,
            xvar,
            yvar,
            xlab = "x",
            ylab = "y",
            groupvar = "Group",
            title = "correlation",
            plot.width = 2.5,
            save = T,
            leg = F,
            xlim = F,
            ylim = F,
            marginal = F) {
    data <- as.data.frame(data)
    
    if (save == T) {
      pdf(
        paste0(today, filename),
        width = plot.width,
        height = plot.width * 1.05,
        pointsize = 10,
        family = 'Helvetica',
        useDingbats = FALSE
      )
    }
    par(
        mar = c(2.3, 2.3, 1, 0.2),
        mgp = c(1.25, 0.5, 0),
        cex.lab = 0.8,
        cex.axis = 0.7,
        cex.main = 0.8,
        tcl = -.35
      )
    
    v1 <- data[, xvar]
    v2 <- data[, yvar]
    g <- data[, groupvar]
    
    xlim <- if (all(xlim == F))
      xlim <- range(v1, na.rm = T)
    ylim <- if (all(ylim == F))
      ylim <- range(v2, na.rm = T)
    
    if(marginal){ 
      par(fig=c(0,0.85,0,0.85),
          mar = c(2.3, 2.3, 0, 0)
          )
      title=NA
      }
    
    plot(
      v2 ~ v1,
      col = pal[g],
      pch = 20,
      xlab = xlab,
      ylab = ylab,
      main = title,
      xlim = xlim,
      ylim = ylim
    )
    m <- lm(v2 ~ v1)
    ct <- cor.test(v2, v1, method = "pearson")
    
    if (ct$p.value < 0.05) {
      abline(m, lwd = 1.5)
      
      newx <-
        seq(min(v1, na.rm = T) * 0.9,
            max(v1, na.rm = T) * 1.1,
            length.out = length(v1))
      prd <-
        predict.lm(
          m,
          newdata = data.frame(v1 = newx),
          interval = c("confidence"),
          level = 0.90,
          type = "response"
        )
      lines(newx, prd[, 2], col = "grey", lty = 2)
      lines(newx, prd[, 3], col = "grey", lty = 2)
    }
    mtext(
      paste(c(
        "r =",
        round(unlist(ct[c("estimate")]), digits = 2),
        ", p =",
        signif(ct$p.value, digits = -1)
      ), collapse = ""),
      side = 3,
      cex = 0.8,
      line = -1.2
    )
    if (leg == T)
      legend(
        "bottomright",
        levels(g),
        pch = 16,
        cex = 0.8,
        col = pal[factor(levels(g), levels = levels(g))],
        inset = 0.02
      )
    # Add marginal boxplots to the scatterplot
    if(marginal){
      
      par(fig=c(0,0.85,0.86,1), 
          mar=c(0, 2.3, 0.1, 0), new=TRUE)
      plot.bxp("filename", data, xvar, ylab1=NA,
                saveit = F,
                marginal = T,
                horizontal=T)
      
      par(fig=c(0.86,1,0,0.85), 
          mar = c(2.3, 0, 0, 0.1),
          new=TRUE
          )
      
      plot.bxp("filename", data, yvar, ylab1=NA,
               saveit = F,
               marginal = T)
      
      mtext(title, cex=0.7, side=3, outer=TRUE, line=-5)
    }
    
    
    if (save == T) dev.off()
  }

#### Read files and subset useful data ####
setwd("~/UPGON/LIN-5/MANUSCRIPT/Reviews/Figures/Figure 5") # change to the source data location on your machine
dt <- read.xlsx("Figure 5 - source data.xlsx")

# factor for upshifted dead, upshifted hatched, controls...
dt$Group <-
  factor(dt$Group, levels = c("Ctrl", "Alive", "Dead", "Inverted"))

equal <- dt$AB.rel > 0.48 & dt$AB.rel <= 0.53
ctrl <- grepl("JT.", dt$ID)
inv <- dt$AB.rel <= 0.48

#keep only equalized embryos >  omit partially equalized and inverted embryos
dt <- dt[inv | equal | ctrl, ]

#### E2 division delay after MSp ####
dt$Ea.LifeTime <- (dt$Ea - dt$E)
dt$Ep.LifeTime <- (dt$Ep - dt$E)

dt$MS.duration <- dt$MSp - dt$MS # effectively cell cycle duration of MSp cell
dt$E2.duration <- (dt$Ea.LifeTime + dt$Ep.LifeTime) / 2

dt$Ea.delay <- dt$Ea - dt$MSp
dt$Ep.delay <- dt$Ep - dt$MSp
dt$E2.delay <- (dt$Ea.delay + dt$Ep.delay) / 2

# Figure 5C ####
plot.bxp(
  filename = "_E2_delay_after.MSp.pdf",
  data = dt,
  saveit = T,
  var1 = "E2.delay",
  ylab1 = "E2 delay after MSp division [min]",
  plot.width = 1
)
compare_means(
  E2.delay ~ Group,
  data = dt,
  method = 't.test',
  p.adjust.method = "BH"
)
aggregate(dt$E2.delay, by = list(dt$Group), mean)
aggregate(dt$E2.delay, by = list(dt$Group), sd)
aggregate(dt$E2.delay, by = list(dt$Group), length)

# Ea and Ep delays individually (not shown in the manuscript)
compare.bxp(
  filename = "_Ea_Ep_delay_after_MSp.pdf",
  data = dt,
  var1 = "Ea.delay",
  var2 = "Ep.delay",
  ylab1 = "Ea division delay after MSp [min]",
  ylab2 = "Ep division delay after MSp [min]"
)

compare_means(
  Ea.delay ~ Group,
  data = dt,
  method = 't.test',
  p.adjust.method = "BH"
)
compare_means(
  Ep.delay ~ Group,
  data = dt,
  method = 't.test',
  p.adjust.method = "BH"
)

plot.corr(
  filename = "_Ea_Ep_delay_corr_equalized.pdf",
  data = dt,
  groupvar = "Group",
  xvar = "Ea.delay",
  yvar = "Ep.delay",
  title = "Division delay after MSp",
  xlab = "Ea [min]",
  ylab = "Ep [min]",
  save = T,
  leg = T
)

# Fig. 5 - supplement - C - marginal boxplots - E2 avg cell cycle duration ####
plot.bxp(
  filename = "_E2_avg_cell_cycle.pdf",
  data = dt,
  var1 = "E2.duration",
  ylab1 = "mean cell cycle duration of Ea/Ep [min]",
  plot.width = 1
)

compare_means(
  E2.duration ~ Group,
  data = dt,
  method = 't.test',
  p.adjust.method = "BH"
)
# not signif difference betwwen alive and dead, but very different distribution
  
  aggregate(dt$E2.duration,by = list(dt$Group),mean,na.rm = T)
  aggregate(dt$E2.duration, by = list(dt$Group), sd, na.rm = T)
  aggregate(dt$E2.duration, by = list(dt$Group), length)

# Ea, Ep Cell cycle duration (separately, not shown)

compare.bxp(
  filename = "_E2_cell_cycle_duration.pdf",
  data = dt,
  var1 = "Ea.LifeTime",
  var2 = "Ep.LifeTime",
  ylab1 = "Ea cell cycle duration [min]",
  ylab2 = "Ep cell cycle duration [min]",
  plot.width = 2
)

compare_means(
  Ea.LifeTime ~ Group,
  data = dt,
  method = 't.test',
  p.adjust.method = "BH"
)
compare_means(
  Ep.LifeTime ~ Group,
  data = dt,
  method = 't.test',
  p.adjust.method = "BH"
)

#Ingression ####
dt$E2.disp <- (dt$EaDisp + dt$EpDisp) / 2
# Fig. 5D ####
plot.corr(
  filename = "_E2_ingression~AB.size_corr.pdf",
  data = dt,
  groupvar = "Group",
  xvar = "AB.rel",
  yvar = "E2.disp",
  title = "E2 trajectory  ~ AB size %",
  xlab = "AB size %",
  ylab = "Ea/Ep trajectory [um]"
)

# Fig. 5D - marginal boxplots - E2 ingression - trajectory length during Ea/Ep cell cycle ####
plot.bxp(
  filename = "_E2_ingression_distance.pdf",
  data = dt,
  var1 = "E2.disp",
  ylab1 = "E2 displacement [um]",
  plot.width = 1
)
compare_means(
  E2.disp ~ Group,
  data = dt,
  method = 't.test',
  p.adjust.method = "BH"
)
aggregate(dt$E2.disp, by = list(dt$Group), mean, na.rm = T)
aggregate(dt$E2.disp, by = list(dt$Group), sd, na.rm = T)
aggregate(dt$E2.disp, by = list(dt$Group), length)

# Fig. 5 - supplement - A -  migration of Ea and Ep indvidually ####
compare.bxp(
  filename = "_Ea_Ep_ingression.pdf",
  data = dt,
  var1 = "EaDisp",
  var2 = "EpDisp",
  ylab1 = "Ea migration [um]",
  ylab2 = "Ep migration [um]"
)
compare_means(EaDisp ~ Group, data = dt, method = 't.test', p.adjust.method = "BH") #NS
compare_means(EpDisp ~ Group, data = dt, method = 't.test', p.adjust.method = "BH") #NS

# Fig. 5 - supplement - B ####
plot.corr(
  filename = "_E2_ingression~E2.CellCycle_corr.pdf",
  data = dt,
  groupvar = "Group",
  xvar = "E2.duration",
  yvar = "E2.disp",
  title = "E2 trajectory  ~ Cell cycle",
  xlab = "E2 cell cycle [min]",
  ylab = "E2 trajectory [um]"
)

#correlation between Ea and Ep (not shown)
plot.corr(
  filename = "_Ea_Ep_ingression_corr.pdf",
  data = dt,
  groupvar = "Group",
  xvar = "EaDisp",
  yvar = "EpDisp",
  title = "Ingression distance",
  ylab = "Ea trajectory [um]",
  xlab = "Ep trajectory [um]"
) # not well correlated (ns)

# END-3::GFP expression ####
dt$E2exp <- (dt$EaMax+dt$EpMax)/2
  
# Fig. 5F ####
#correlation of AB size with mean expression of END-3::GFP in E2 cells
plot.corr(
  filename = "_END-3_expression~AB.size_corr.pdf",
  data = dt,
  groupvar = "Group",
  xvar = "AB.rel",
  yvar = "E2exp",
  title = NA,
  xlab = "AB size %",
  ylab = "Ea/Ep END-3::GFP expression [A.U.]",
  marginal=T
)


#5F marginal boxplots ####
plot.bxp(
  filename = "_END-3-GFP_expression.pdf",
  data = dt,
  saveit = T,
  var1 = "E2exp",
  ylab = "Ea/Ep END-3::GFP expression [A.U.]",
  plot.width = 1
)

plot.bxp(
  filename = "_AB.size.pdf",
  data = dt,
  saveit = T,
  var1 = "AB.rel",
  ylab = "Ea/Ep END-3::GFP expression [A.U.]",
  plot.width = 1
)

compare_means(E2exp ~ Group, data = dt, method = 't.test', p.adjust.method = "BH")
aggregate(dt$E2exp, by = list(dt$Group), mean)
aggregate(dt$E2exp, by = list(dt$Group), sd)
aggregate(dt$E2exp, by = list(dt$Group), length)

aggregate(dt$E2exp, by = list(dt$Proc), mean)
aggregate(dt$E2exp, by = list(dt$Proc), sd)

t.test(E2exp~Proc, data=dt)#controls vs equalized expression


#Association between Elt-2::GFP and outcome - only in equalized ####
#          alive	dead
# normal	    2	    3
# abnormal	  4	    7
outcomes <- matrix(c(2, 4, 3, 7), nrow=2, ncol=2, dimnames=list(c("normal","abnormal"),c("Alive","Dead")))
fisher.test(outcomes) #p-value = 1 >> compression leads to elevated lethality

#Association between Elt-2::GFP and outcome - all including inverted
#          alive	dead
# normal	  5	    5
# abnormal	6	    10
outcomes <- matrix(c(5, 6, 5, 10), nrow=2, ncol=2, dimnames=list(c("normal","abnormal"),c("Alive","Dead")))
fisher.test(outcomes) #p-value = 0.689 >> compression leads to elevated lethality

# Fig. 5 - supplement C ####
plot.corr(
  filename = "_END-3_expression~E2_cycle_corr.pdf",
  data = dt,
  groupvar = "Group",
  xvar = "E2.duration",
  yvar = "E2exp",
  title = NA,
  xlab = "Ea/Ep cell cycle [min]",
  ylab = "Ea/Ep END-3::GFP expression [A.U.]"
)
```
